# Transcriptomes of plant gametophytes have a higher proportion of rapidly evolving and young genes than sporophytes

**DOI:** 10.1101/022939

**Authors:** Toni I. Gossmann, Dounia Saleh, Marc W. Schmid, Michael A. Spence, Karl J. Schmid

**Author notes:** Co-First Authors (Toni I. Gossmann, Dounia Saleh). Current address: S3IT – Service and Support for Science IT, University of Zurich, Switzerland.

## Abstract

Reproductive traits in plants tend to evolve rapidly due to various causes that include plant-pollinator coevolution and pollen competition, but the genomic basis of reproductive trait evolution is still largely unknown. To characterise evolutionary patterns of genome wide gene expression in reproductive tissues in the gametophyte and to compare them to developmental stages of the sporophyte, we analysed evolutionary conservation and genetic diversity of protein-coding genes using microarray-based transcriptome data from three plant species, *Arabidopsis thaliana*, rice (*Oryza sativa*) and soybean (*Glycine max*). In all three species a significant shift in gene expression occurs during gametogenesis in which genes of younger evolutionary age and higher genetic diversity contribute significantly more to the transcriptome than in other stages. We refer to this phenomenon as “evolutionary bulge” during plant reproductive development because it differentiates the gametophyte from the sporophyte. We show that multiple, not mutually exclusive, causes may explain the bulge pattern, most prominently reduced tissue complexity of the gametophyte, a varying extent of selection on reproductive traits during gametogenesis as well as differences between male and female tissues. This highlights the importance of plant reproduction for understanding evolutionary forces determining the relationship of genomic and phenotypic variation in plants.

## Introduction

Reproductive traits in plants and animals tend to be highly diverse and rapidly evolving within and between closely related species (Swanson and Vacquier, 2002; Barrett, 2002; Parsch and Ellegren, 2013). Their diversity may be influenced by the coevolution with pollinators or pathogens that infect reproductive tissues, the mating system (i.e. selection for the maintenance of self-incompatibility), the rapid evolutionary dynamics of sex chromosomes, genomic conflicts between parents and offspring, or from sexual selection (Baack et al., 2015). Some genes and proteins expressed in reproductive tissues exhibit high rates of evolution (Swanson and Vacquier, 2002; Parsch and Ellegren, 2013). In plants, they include genes encoding the self-incompatibility system (Nasrallah et al., 2002; Tang et al., 2007), pollen-coat proteins (Schein et al., 2004) and imprinted genes controlling resource allocation to offspring (Spillane et al., 2007). The rapid evolution of reproductive traits and their underlying genes is in contrast to other tissues and developmental stages that appear to be more conserved. In particular, the phylotypic stage in animals, in which a similar morphology at a certain stage of embryo development is observed within phyla, represents the archetype of morphological evolutionary conservation within a phylum (Duboule, 1994).

Although reproductive traits appear to evolve rapidly in animals, plants and other organisms with anisogamic sexual reproduction (Lipinska et al., 2015), there is a fundamental difference between these groups. In animals, a group of cells are set aside during early development, which forms the germ line. Plants do not have a germ line, but are characterized by alternating sporophytic and haploid gametophytic stages (Schmidt et al., 2011; Grossniklaus, 2011). Since the two stages differ in their development and role in reproduction, the function and evolution of genes expressed in the sporophyte and gametophyte should also differ. Furthermore, the haploid stage immediately exposes recessive mutations to selection which causes different evolutionary dynamics of genes expressed in the gametophyte compared to genes only expressed in a diploid stage (Gossmann et al., 2014b).

Currently it is little understood which processes drive the rapid evolution of plant reproductive genes on a genome-wide scale. During plant gametogenesis, the transcription profile changes dramatically, and genes involved in reproduction are enriched in this phase (Schmid et al., 2005; Fujita et al., 2010; Xiao et al., 2011; O’Donoghue et al., 2013). However, a focus on genes whose expression is enriched in a specific tissue introduces a bias for genes with specific expression patterns that ignores the contribution of other genes to the total diversity of expression patterns (Arunkumar et al., 2013; Gossmann et al., 2014b). To characterise the evolutionary dynamics of transcriptomic profiles it is therefore necessary to combine the genome-wide expression intensity of all genes expressed in a given tissue and stage with evolutionary parameters quantifying the level of polymorphism, rate of molecular evolution or long-term evolutionary conservation (Slotte et al., 2011). For this purpose, evolutionary indices such as the transcriptome age index (TAI), which measures the long-term conservation of expressed genes weighted by the relative expression of the gene, or the divergence index (TDI), which compares the rate of non-synonymous to synonymous substitutions in a proteincoding gene between closely related species (Domazet-Lošo and Tautz, 2010; Kalinka et al., 2010; a molecular equivalent. Studies in vertebrates (zebrafish) and insects (*Drosophila melanogaster*) confirmed this hypothesis because genes expressed during the phylotypic stage were more conserved and less rapidly evolving than genes expressed in other stages of development (Domazet-Lošo and Tautz, 2010; Kalinka et al., 2010). Although plants do not have a clear morphologically defined phylotypic stage, a transcriptomic hourglass was also postulated for the model plant *Arabidopsis thaliana* because old and slowly evolving genes contribute disproportionally to the overall transcriptome during early stages of embryo development (Quint et al., 2012; Drost et al., 2015), but see Piasecka et al. (2013).

Based on the above considerations, we reasoned that the morphologically and developmentally diverse reproductive stages of plants, in particular the gametophyte, should be characterized by a high proportion of expressed genes with a lower degree of long-term evolutionary conservation (Cui et al., 2015) and a higher rate of divergence between closely related species. We tested this hypothesis by comparing the transcriptome-based indices of evolution observed in reproductive stages like the gametogenenesis to other developmental stages such as the putative phylotypic stage. We based our analysis on three different evolutionary parameters and used gene expression and genome sequence data from three flowering plant species, *Arabidopsis thaliana*, rice (*Oryza sativa*), soybean (*Glycine max*), and the moss *Physcomitrella patens*. The expression data include developmental stages preceeding (e.g. flower development), during and following gametogenesis (e.g. embryogenesis). The *A. thaliana* data additionally included stages from both sexes, while for the other species we used data from the male sex only. Our results show that the rate of evolution of genes expressed in reproductive stages is much higher relative to the extent of conservation of the putative phylotypic or other sporophytic stages. For this reason, we name this observation ‘evolutionary bulge’ to express the stronger contribution of rapidly evolving and young genes to the transcriptome in reproductive developmental stages compared to other stages and discuss several, not mutually exclusive, hypotheses that may explain this pattern.

## Results and Discussion

To test whether developmental stages and tissues involved in reproduction show a higher proportion of expressed genes of a younger evolutionary age and a higher rate of divergence between closely related species, we analysed global expression during gamete development and the developmental stages before and after gametogenesis (Table 1) with three evolutionary parameters. For this we combined microarray expression levels with measures of evolutionary conservation and polymorphism into evolutionary transcriptome indices of developmental stages. The evolutionary transcriptome index is calculated as:

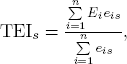

**Table 1:** Summary of microarray-based expression data from different developmental stages used in this study. Further details about the individual datasets are provided in Supporting File S1.

| Species | Developmental Stage | References |
| --- | --- | --- |
| *A. thaliana* | *Pre-Reproductive stage:* Shoot apex 7 days (SA7D), Shoot apex 14 days (SA14D), Shoot after bolting (SAB), Flower stage 9 (FS9), Flower stage 12 (FS12), Flower stage 15 (FS15)<br>*Reproductive stage:* Megaspore mother cell (MMC), Egg cell (EC), Unicellular pollen (UCP), Bicellular pollen (BCP), Tricellular pollen (TCP), Pollen mature (MP), Sperm (S), Pollentube (PT)<br>*Post-reproductive stage:* Quadrant embryo (Q), Globular embryo (G), Heart embryo (H), Torpedo embryo (T), Mature embryo (M) | Schmid et al. (2005)<br>Honys and Twell (2004); Borges et al. (2008); Wang et al. (2008); Wuest et al. (2010); Schmidt et al. (2011); Schmid et al. (2012)<br>Le et al. (2010); Zuber et al. (2010) |
| Rice | *Pre-Reproductive stage*<br>Shoot 4 weeks (S4W)<br>*Reproductive stage:* Unicellular pollen (UCP), Bicellular pollen (BCP), Tricellular pollen (TCP), Mature pollen (MP), Germinated pollen (GP)<br>*Post-Reproductive stage:* Fertilisation (F), Zygote formation (Z), 0 Days After Pollination embryo (0DAP), 1 Days After Pollination embryo (1DAP), 2DAP embryo, 3DAP embryo, 4DAP embryo, 9DAP embryo, 12DAP embryo | Fujita et al. (2010)<br>Wei et al. (2010)<br>Fujita et al. (2010); Gao and Xue (2012) |
| Soybean | *Pre-Reproductive stage:* Sporophyte (S)<br>*Reproductive stage:* Mature pollen (MP)<br>*Post-Reproductive stage:* Globular embryo (G), Heart embryo (H), Cotyledon (C), Seed parenchym (SP), Seed meristem (SSM) | Haerizadeh et al. (2009)<br>Haerizadeh et al. (2009)<br>Le et al. (2007) |

where E is the evolutionary parameter, *s* the developmental stage, *E_i_* the value of the evolutionary parameter for gene *i, n* the total number of genes and *e_is_* the expression level of gene *i* in developmental stage *s*. In this study, we used gene age to calculate the transcriptomic age index (TAI) (Kalinka et al., 2010; Domazet-Lošo and Tautz, 2010), sequence divergence (*d_N_* / *d_S_*) for the transcriptomic divergence index (TDI) and sequence diversity (*p_N_* / *p_S_*) for new transcriptome polymorphism index (TPI), which is a measure of current evolutionary constraint. The evolutionary transcriptome index is related to Pearson’s correlation coefficient but also incorporates variation in expression mean and variation (Supplementary text S1). This statistic is different from previous approaches addressing similar questions of evolutionary patterns during reproduction. Instead of focusing on significantly enriched genes which are biased towards specifically and/or strongly expressed genes, we considered the composition of the whole transcriptome. This enabled us to differentiate whether any evolutionary signals during development are caused by a few genes with strong effects or many genes with weak effects. It also allows to directly compare signal intensities with the previously described evolutionary hourglass during embryo development in *A. thaliana*.

In all three species we observed the highest values of the three indices during reproductive stages (Figure 1), and they differ significantly from the values of the sporophytic developmental stages. To exclude that high point estimates of evolutionary parameters, which may be caused by low quality alignments, inflate diversity and polymorphism indices, we calculated TDI and TPI values from the weighted median (see Material and Methods). Both indices are robust to the impact of low quality alignments of few genes (Supplementary Figure S1). Large absolute differences in the expression level of genes with a high and low expression level may allow a few genes to dominate the overall transcriptome index. We conducted our analyses with *log*_2_ transformed data, but additionally verified the bulge pattern with raw and *log*_10_-transformed expression data and found that the transcriptome indices are little influenced by genes with very high expression levels (Supplementary Figure S2). In *A. thaliana*, pollen tubes have the highest TAI value and therefore the highest proportion of young genes (*t*-test; *P* < 6.5 × 10^-34^ for all pairwise comparisons with sporophytic stages). The highest TDI and TPI values occur in sperm cells (*P* < 2.2 × 10^-15^). In rice, the highest TAI, TDI and TPI indices are observed in the mature and germinated pollen stages (*P* < 6 × 10^-27^ for all pairwise comparisons), and in soybean in the germinated pollen stage (*P* < 7.3 × 10^-6^). The *A. thaliana* and rice expression data cover consecutive reproductive stages in which the evolutionary indices increase during the maturation of the male gametes and peak at a final reproductive stage. Female gametophytic tissues show a similar trend in *A. thaliana*. Overall, there is a strong difference between gametophytic and sporophytic phases, suggesting a distinct evolutionary dynamic of reproductive compared to sporophytic stages. The comparison of evolutionary indices between pre- and postgametic developmental stages reveal that the lowest values of these indices are not consistently the lowest during embryogenesis, as suggested by the hourglass hypothesis. Except for *A. thaliana*, there is no particular stage during embryogenesis that has the lowest TAI, TDI and TPI values (Figure 1).

**Figure 1:**
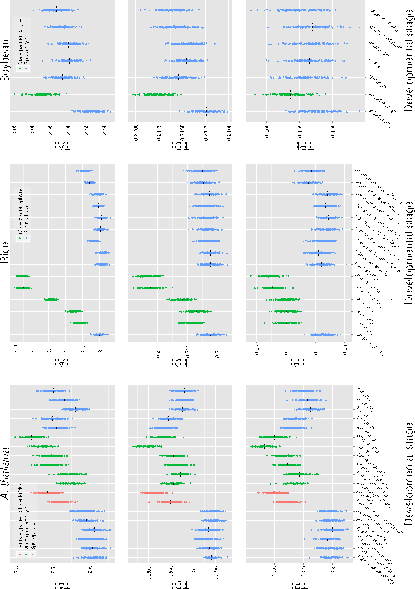
Evolutionary transcriptome indices for *A. thaliana*, rice and soybean. Plot of transcriptome age index (TAI), transcriptome divergence index (TDI) transcriptome polymorphism index (TPI) for available data from *Arabidopsis thaliana*, rice and soybean for different developmental stages and tissues. Black lines indicate the transcriptome index and the coloured dots are the indices calculated from

All transcriptome data for a given species were generated with the same Affymetrix array, but hybridisations were conducted in independent experiments. To test for confounding effects from the experimental conditions we also calculated the transcriptome indices by pre-processing datasets independently (Supplementary Figure S3). This led to a relative shift of transcriptome indices between pre- and postgametophytic developmental stages, but the evolutionary bulge remained as a robust pattern. Using *P*-values associated with gene expression from a larger dataset for *A. thaliana* (Supplementary Table S1) we calculated modified transcriptome indices (see Methods) by including only genes that are significantly expressed in a given stage with an FDR < 0.1 (Supplementary Figure S4). With few exceptions, reproductive tissues have higher evolutionary indices, and the number of significantly expressed genes differs between the reproductive and vegetative phase (Pina et al., 2005) (*P* = 2 × 10^-12^, U-test of the median number of genes significantly expressed in reproductive versus sporophytic tissues).

Since the three evolutionary indices may not be independent of each other, we analysed their correlation with expression and accounted for potentially co-varying factors (Gossmann et al., 2014a). By assuming that expression variation between samples is similar and the same genes are analysed across stages, the evolutionary index is proportional to the correlation coefficient, *r* (For a derivation, see Supplementary Text S1). The analysis of correlation supports the evolutionary bulge pattern because the highest value of *r* is observed for the gametophytic stages (Table 2; subset of sporophytic and gametophytic stages). The only exception was the polymorphism index (TPI) of the two domesticated species (rice and soybean) which was influenced in the reproductive stage by differences in expression variance between reproductive and sporophytic stages (Supplementary Figure S5). Results of partial correlations, taking the other two evolutionary parameters, as well as gene length and *d_S_* (a proxy for mutation rate) as co-variates, are qualitatively very similar to the pairwise correlations (Table 2). Patterns of polymorphism in domesticated species are affected by past domestication bottlenecks (Gossmann et al., 2010) and the global expression pattern of domesticated species may be substantially altered (e.g., Rapp et al., 2010; Yoo and Wendel, 2014). Since the evolutionary bulge pattern is influenced by different processes in the three species (Figure 2 and Supplementary Figure S6), domestication may explain some differences of TPI values between the wild and the two crop plant species.

**Table 2.**
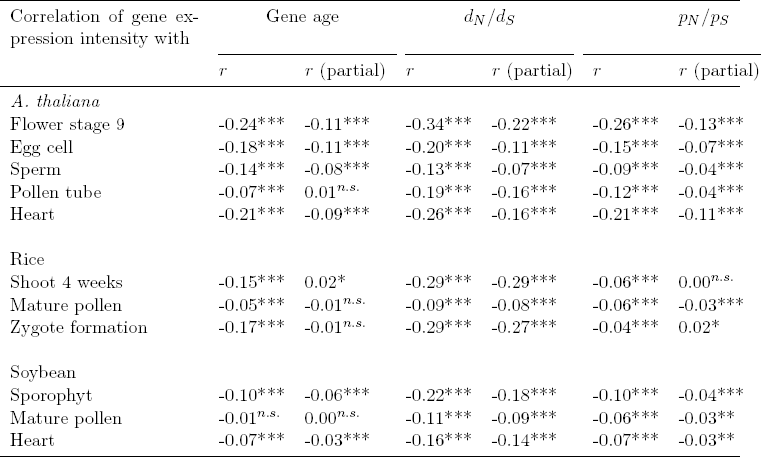
Correlation of gene expression with three evolutionary indices. The analysis was based on Pearson’s correlation and partial correlation for selected development stages. For the partial correlations, the other two evolutionary parameters as well as gene length and *d_s_* were used as co-variates.

| Correlation of gene expression intensity with | Gene age | | $d_N/d_S$ | | $p_N/p_S$ | |
| --- | --- | --- | --- | --- | --- | --- |
| | $r$ | $r$ (partial) | $r$ | $r$ (partial) | $r$ | $r$ (partial) |
| *A. thaliana* |  |  |  |  |  |  |
| Flower stage 9 | -0.24*** | -0.11*** | -0.34*** | -0.22*** | -0.26*** | -0.13*** |
| Egg cell | -0.18*** | -0.11*** | -0.20*** | -0.11*** | -0.15*** | -0.07*** |
| Sperm | -0.14*** | -0.08*** | -0.13*** | -0.07*** | -0.09*** | -0.04*** |
| Pollen tube | -0.07*** | 0.01<sup>n.s.</sup> | -0.19*** | -0.16*** | -0.12*** | -0.04*** |
| Heart | -0.21*** | -0.09*** | -0.26*** | -0.16*** | -0.21*** | -0.11*** |
| Rice |  |  |  |  |  |  |
| Shoot 4 weeks | -0.15*** | 0.02* | -0.29*** | -0.29*** | -0.06*** | 0.00<sup>n.s.</sup> |
| Mature pollen | -0.05*** | -0.01<sup>n.s.</sup> | -0.09*** | -0.08*** | -0.06*** | -0.03*** |
| Zygote formation | -0.17*** | -0.01<sup>n.s.</sup> | -0.29*** | -0.27*** | -0.04*** | 0.02* |
| Soybean |  |  |  |  |  |  |
| Sporophyt | -0.10*** | -0.06*** | -0.22*** | -0.18*** | -0.10*** | -0.04*** |
| Mature pollen | -0.01<sup>n.s.</sup> | 0.00<sup>n.s.</sup> | -0.11*** | -0.09*** | -0.06*** | -0.03** |
| Heart | -0.07*** | -0.03*** | -0.16*** | -0.14*** | -0.07*** | -0.03** |

**Figure 2:**
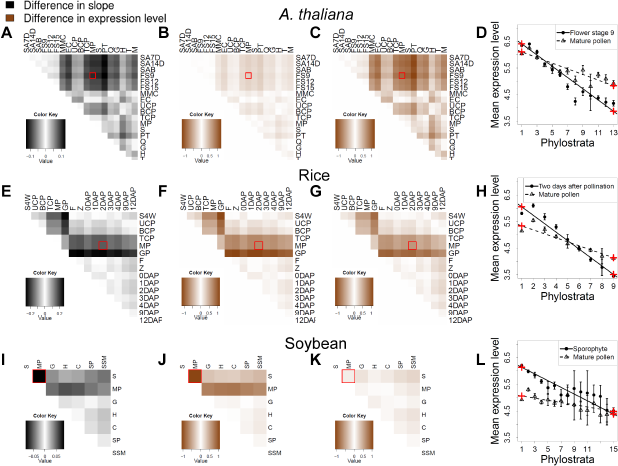
Difference in expression level between young and old genes and between developmental stages. (a-d) *A. thaliana* (e-h) rice (i-l) soybean (a, e, i) Heatmaps of differences in linear regression slopes between pairs of developmental stages included in the analysis. (b, f, j) Heatmaps of differences in expression level inferred from linear regressions between pairs of developmental stages for the first phylostratum (PS= 1). (c, g, k) Heatmaps of differences in expression level inferred from linear regressions between pair of developmental stages for the youngest phylotratum (PS= 13 in *A. thaliana*; PS= 9 in rice; and PS= 15 in soybean). (d, h, l) Mean, confidence interval and linear regression of expression level for several phylostrata at two stages: Flower stage 9 and mature pollen in *A. thaliana*, 2DAP and mature pollen in rice, sporophyte and mature pollen in soybean. Red crosses represent the expression level inferred from the linear regressions for PS=1 and PS=13/9/15, respectively. For abbreviations of developmental stages, see Supplementary Table S1.

Different expression patterns during gamete development may result from up-regulation of young or down-regulation of old genes and may cause the bulge pattern. We performed linear regression of mean *log*_2_ normalized expression intensities over the gene age of each stage (Figure 2) to infer how strongly the correlation varied between stages. To illustrate changes in expression for different gene ages we selected a pairwise comparison between mature pollen and a sporophytic stage for each species as an example (Figure 2). In all three species, the relative expression of both old and young genes differed between developmental stages, but the extent of change varied between stages and species. In *A. thaliana*, the differences were mainly caused by a change in the expression level of young genes (Figure 2b and c) and in rice by a higher expression of young and a lower expression of older genes (Figure 2f and g). In soybean, the change in expression was mainly caused by the lower expression level of old genes (Figure 2j and k). We also compared the expression levels between stages by grouping genes by their average values of *d_N_*/ *d_S_* and *p_N_* / *p_S_* (Supplementary Figure S6) to test whether expression levels differ between slow and rapidly evolving genes. In *A. thaliana*, conserved genes (low *d_N_* / *d_S_* and *p_N_*/*p_S_*) showed a lower expression level and divergent genes (high *d_N_* / *d_S_* and *p_N_*/*p_S_*) a higher expression level in reproductive stages, especially in pollen and pollen tubes. In rice, genes with low *d_N_*/*d_S_* and *p_N_*/*p_S_* values showed strongly decreased mean expression levels in reproductive stages, whereas in soybean, mean expression levels decreased independently from d_N_/d_S_ and p_N_/p_S_ during reproduction.

During reproductive development the tissue complexity of the gametophyte in higher plants is reduced to single cells or a few cells suggesting a reduced interaction between cells and cell types compared to other stages. Highly connected genes tend to evolve slower as a consequence of their functional importance (Alvarez-Ponce and Fares, 2012). Such genes, however, may be less expressed in the gametophytic stage and therefore contribute less to the bulge pattern. This hypothesis is supported by a reduced expression level of old genes in all three species (Figure 2b, f, j). Using data from the *Arabidopsis* interactome database (see Methods) we found that in the late stages of male gametophytes the level of interactions is reduced and shows the lowest value in the pollen tube (Figure 3, *P* < 0.03). In the female gametophyte, which is a tissue of higher complexity, such a reduction in protein interactions is not observed. This difference suggests that factors contributing to the evolutionary bulge pattern may vary between male and female tissues.

**Figure 3:**
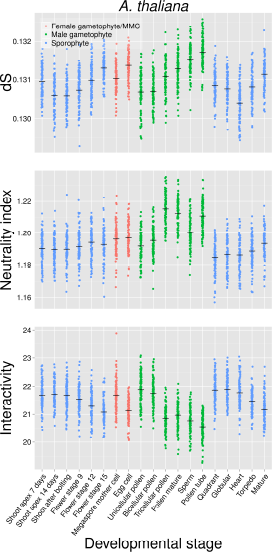
Transcriptome indices for *d_S_*, neutrality index and gene interactions for *A. thaliana*. Upper panel: Median per gene *d_S_* (synonymous per site substitution rate, a proxy for the neutral mutation rate) weighted by gene expression. Middle panel: Median per gene neutrality index (NI, a measurement of the departure from neutrality, with NI≈ 1 indicating neutrality) weighted by gene expression. Lower panel: Average number of gene interaction partners weighted by gene expression.

An evolutionary bulge pattern might be relatively less pronounced in self-fertilizing species, like the three species analysed here, as they lack genetic diversity (Wright et al., 2013) and deleterious recessive mutations are rapidly removed in diploid tissues (Szövényi et al., 2014). On the other hand, an evolutionary bulge pattern should be independent from the mating system if low but sufficient levels of outcrossing occur in selfers (Bomblies et al., 2010), if most mutations are dominant and therefore exposed to selection in outcrossers, or if the reproductive success of the gametophyte is dominated by *de novo* mutations during gametogenesis. The silent sequence divergence between species, *d_S_*, is a proxy for mutation rate and is increased for genes predominantly expressed in sperm and pollen tube stages in *A. thaliana* (Figure 3; *P* < 1.7 × 10^-4^) which supports the latter explanation.

Mosses have an extended generation of multicellular haploid gametophytes that differentiate into early vegetative and later reproductive stages and allow to investigate the effects of haploidy on transcriptome indices. In the expression data available for gametophytic and sporophytic stages of the moss *Physcomitrella patens* (O’Donoghue et al., 2013), young genes contribute to the gene age of the gametophytic transcriptome as indicated by an increase of the TAI during the haploid stage (Figure 4; *P* < 3.2 × 10^-10^). This is consistent with the evolutionary bulge and suggests that it may be a general pattern of plant reproductive evolution, although a broader taxonomic sampling will be necessary to verify this hypothesis.

**Figure 4:**
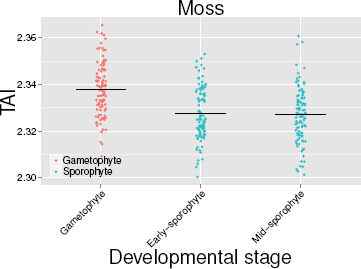
Estimates of the transcriptomic age index (TAI) for three different developmental stages in the moss *Physcomitrella patens*.

The pollen tube of *A. thaliana* showed lower TDI and TPI, but higher TAI values than the sperm cell (Figure 1; see also Cui et al., 2015), which indicates that tissue- or cell-specific effects weighted neutrality index (NI; NI< 1 indicates an increased role of positive selection while NI> 1 indicates purifying selection) differs between sperm and late pollen stages in *A. thaliana* (Figure 3, *P* < 2.7 × 10^-13^) which shows a shift in the relative contribution of positive and negative selection and supports tissue-specific effects. A possible explanation is an enrichment of slightly deleterious mutations that are more effectively removed in pollen due to purifying selection, but it is difficult to disentangle the extent of the different selective forces on a gene-by-gene basis. As noted before, a focus on tissue-specific enriched genes represents a bias because these genes tend to show a narrow expression pattern and a high expression level. In plants, both factors correlate with the rate of molecular evolution, but in opposite directions (Slotte et al., 2011).

## Conclusion

When compared to the transcriptomic hourglass of embryogenesis, the evolutionary bulge seems to be a more robust pattern of plant development. We reproduced the hourglass in *A. thaliana*, but found little support for it in rice or soybean which may result from an incomplete sampling of embryonic stages in the latter two species. This suggests that the hourglass pattern is restricted to a very short time span of plant embryo development. Therefore, further research is required to verify the transcriptomic hourglass as a general pattern of plant development because the transcriptome indices are not consistently lower during embryogenesis than in other developmental stages. In contrast, the evolutionary bulge of reproduction is seen in four plant species illustrating that the evolutionary forces acting during plant reproductive development leave a strong imprint on the genomic composition of protein-coding genes. This is consistent with the phenotypic diversity of reproductive traits but additionally highlights the importance of plant reproduction for understanding evolutionary forces determining the relationship of genomic and phenotypic variation in plants. We have shown that multiple, not mutually exclusive, causes may explain the bulge pattern, most prominently reduced tissue complexity of the gametophyte and a varying extent of selection on reproductive traits during gametogenesis as well as between male and female tissue. To further test whether the evolutionary bulge is a general pattern of plant evolution and to disentangle the different factors that are influencing it, the investigation of plant species with strong differences in their mode of reproduction in comparison to our study species will be useful. Examples are diecious plants, wind-pollinated outcrossing trees, insect-pollinated flowering plants and species with increased complexity of the gametophyte generation.

## Materials and Methods

### Sequence data and software

We obtained the genome sequences of *A. thaliana* (Arabidopsis Genome Initiative, 2000), rice (*Oryza sativa*, International Rice Genome Sequencing Project 2005) and soybean (*Glycine max*, Schmutz et al. 2010) from the plant genome database (Duvick et al., 2008) and the plant duplication database (Lee et al., 2013) along with their outgroups *Arabidopsis lyrata* (Hu et al., 2011), *Sorghum bicolor* (Paterson et al., 2009) and *Phaseolus vulgaris* (Schmutz et al., 2014), respectively. Polymorphism data were obtained from 80 *Arabidopsis thaliana accessions* (Cao et al., 2011). To identify coding SNP information for rice we used the Rice Haplotype Map Project Database (2nd Generation, http://www.ncgr.ac.cn/RiceHap2/index.html) and soybean we used SNP information deposited in SNPdb (Sherry et al., 2001) and extracted coding SNPs from the soybean genome annotation. We used R and Python scripts to conduct statistical analyses.

### Gene expression data

Gene expression data were obtained for the three plants species from the PlexDB database (Dash et al., 2012) and GEO databases (Barrett et al., 2013). In particular, we focused on development stages preceding gametogenesis, during gametogenesis and embryogenic developments (Table 1 and Supplementary File S1). For each species, Robust Multi-array Analysis (RMA; Irizarry et al., 2003) and invariant set (IS) methods were performed with the affy Bioconductor package to normalize all datasets simultaneously. Scatterplots of expression between replicates showed better results for RMA normalization (data not shown). Therefore, unless stated otherwise, expression data shown in this study are based on a normalisation across experiments using RMA with *log*_2_ transformation. Since different laboratory conditions can affect expression patterns (Massonnet et al., 2010), we controlled for these effects in the *A. thaliana* data (Schmid et al., 2005) by removing datasets that were obtained from plants with different growth conditions before RNA extraction (Supplementary File S1). To check whether the differences in expression between experimental conditions were negligible compared to the differences between stages, we generated scatterplots for the mature pollen stage (Supplementary Figure S7) that was common to different experiments (Honys and Twell, 2004; Schmid et al., 2005; Borges et al., 2008; Wang et al., 2008). Scatterplots showed an expression profile that was similar between experiments with RMA normalization over all experiments and when normalized independently (Supplementary Figures S7 b and c) and also showed more variation between expression levels when compared to non-normalized and IS normalized expression (Supplementary Figure S7 a, d and e). Scatterplots between non-normalized experiments and between IS normalized experiments showed less variation in expression levels, but in general, the correlations between expression levels from different experiments were highly independent from the normalization method. For rice and soybean, all experiments were kept for normalization. Gene expression data for *Physcomitrella patens* for mature gaemtophyte, early and mid sporophyte (O’Donoghue et al., 2013) were downloaded from GEO (GSE32928) and the array and genome annotation (V1.6) was obtained from www.cosmoss.org/physcome_project/wiki/Downloads. In this dataset, two samples per chip are hybridized, each with a different fluorescent dye (green Cy3 and red Cy5). Expression values were averaged across samples.

### Evolutionary parameters

We obtained estimates for TAI (transcriptome age index), TDI (transcriptome divergence index) and TPI (transcriptome polymorphism index) for each developmental stage. A transcriptome index is the average of an evolutionary parameter like gene age (TAI), divergence (TDI) and diversity (TPI) that is weighted by the expression level of each gene. Confidence intervals were obtained by bootstrapping, using 100 sets of genes for each experimental stage. For estimates of gene age we followed the procedure of Quint et al. (2012) which is based on the construction of a phylostratigraphic map. We used one-way BLAST (default parameters) hits against a sets of genomes that are assigned to a certain phylostrata and the BLAST hit to the most distant phylostratum defines the gene age (Albà and Castresana, 2007). The oldest genes have a gene age value of 1 and the highest gene age value was assigned to genes that are specific to a given species (youngest genes). For *A. thaliana* we classified 13 phylostrata, 9 for rice, 15 for soybean and 5 for *Physcomitrella patens*. Altogether we used 40 plant genomes, details about the hierarchical order, the genomes assigned to each phylostratum and number of genes with assigned gene age can be found in Supplementary Figure S8. For each species the largest age category was gene age of value 1.

To calculate a per gene estimate of divergence we calculated *d_N_*/*d_S_* using pairwise alignments of homologous genes identified by INPARANOID from the whole genome comparison with its respective outgroup (Remm et al., 2001; Ostlund et al., 2010). We obtained per gene estimates of *d_N_*/*d_S_* (= *K_a_*/*K_s_*) estimates for genes specific to species pairs with the KaKs_calculator (Zhang et al., 2006). We also introduce a new test statistic, the transcriptomic polymorphism index (TPI).

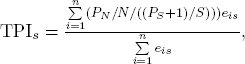

where *s* is the developmental stage, *n* the number of genes, *e_is_* the expression intensity of gene *i* in developmental stage *s*, *P_N_* and *P_S_* the numbers of nonsynonymous and synonymous polymorphisms, respectively, and N and S are the numbers of nonsynonymous and synonymous sites, respectively. We used the ratio of nonsynonymous per site polymorphisms to synonymous per site polymorphism to estimate the distribution of fitness effects. Higher values of *p_N_*/*p_S_* reflect an excess of slightly deleterious mutations (Keightley and Eyre-Walker, 2007). For technical reasons we used *P_S_* + 1 rather than *P_S_* as suggested by Stoletzki and Eyre-Walker (2011) because some genes have no synonymous polymorphisms and therefore would need to be excluded from the analysis which is biased (Stoletzki and Eyre-Walker, 2011). For compactness we refer to the term *P_N_*/*N*/((*P_S_* + 1)/*S*) as *p_N_*/*p_S_* throughout the manuscript.

We tested whether transcriptome indices are different between stages by bootstrapping 100 samples of each index per stage and then performing a two-sample t-test to test for the differences in the means of bootstrapped values. If not noted otherwise, only the highest P-value in the comparison of stages is reported.

### Modified variants of 329 the transcriptome index

We calculated the weighted median transcriptome index of an evolutionary parameter *x* and assumed that 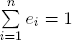. The weighted median of the evolutionary index is then *x_f_* with *f* such that

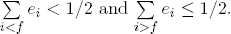

The standardized transcriptome index that does not consider genes with a non-significant expression (Supplementary Figure S4) was calculated as follows:

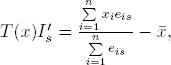

where 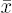 is the arithmetic mean of *x*_1_,…, *x_n_* and *n* the number of significantly expressed genes. We further obtained per gene neutrality index (NI) for *A. thaliana* as follows:

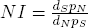

where *p_S_* = (*P_S_* + 1)/*S*. The number of protein interactions for *A. thaliana* were obtained from the *Arabidopsis* interactome database (ftp://ftp.arabidopsis.org/home/tair/Proteins/Protein_interaction_data/Interactome2.0/).

## Acknowledgements

This work was supported by the Deutsche Forschungsgemeinschaft (EVOREP project SCHM1354/7-1) to KJS. The authors are grateful to Arne Jahn, Jonna Kulmuni, Jessica Stapley, two anonymous reviewers and the handling editor for critical comments on the manuscript that have helped to improve the quality of this manuscript.

## Author contributions

TIG and KJS designed the study. TIG, DS and MWS analyzed the data. MAS contributed to analyze the data. TIG and KJS wrote the manuscript. All authors contributed to writing, editing and revising the manuscript.

## Additional information

Supplementary information is available online.

